# High-polyphenol extracts from *Sorghum bicolor* attenuate replication of *Legionella pneumophila* within RAW 264.7 macrophages

**DOI:** 10.1101/2019.12.19.883249

**Authors:** Aubrey M. Gilchrist, Dmitriy Smolensky, Sarah Cox, Ramasamy Perumal, Leela E. Noronha, Stephanie R. Shames

## Abstract

Polyphenols derived from a variety of plants have demonstrated antimicrobial activity against diverse microbial pathogens. *Legionella pneumophila* is an intracellular bacterial pathogen that opportunistically causes a severe inflammatory pneumonia in humans, called Legionnaires’ Disease, via replication within macrophages. Previous studies demonstrated that tea polyphenols attenuate *L. pneumophila* intracellular replication within mouse macrophages via amplification of tumor necrosis factor (TNF) production. *Sorghum bicolor* is a sustainable and resilient cereal crop that thrives in arid environments and is well suited to continued production in warming climates. Polyphenols derived from sorghum have anticancer and antioxidant properties, but their antimicrobial activity has not been evaluated. Here, we investigated the impact of sorghum polyphenols on *L. pneumophila* intracellular replication within RAW 264.7 mouse macrophages. We discovered that sorghum high-polyphenol extract (HPE) treatment attenuates *L. pneumophila* intracellular replication in a dose-dependent manner. Sorghum HPE did not impair bacterial replication *in vitro* or impact macrophage viability. Moreover, in contrast to tea polyphenols, HPE treatment impaired TNF secretion from infected macrophages. Thus, polyphenols derived from sorghum enhance macrophage restriction of *L. pneumophila* by a novel mechanism. This work provides a foundation for the use of sorghum as an antimicrobial agent.

## Introduction

Plant polyphenols are bioactive compounds that naturally function in plant defense against biotic and abiotic stressors. However, polyphenols can also promote human health through antioxidant, anticancer and antimicrobial activity [1–4]. *Camellia sinensis* (tea) polyphenols (catechins) have demonstrated antimicrobial activity against a variety of bacterial, viral and fungal pathogens [5]. Specifically, the tea polyphenol epigallocatechin gallate (EGCg) has broad antimicrobial activity against a variety of pathogens *in vitro* and within eukaryotic host cells [5–8]. With the increased prevalence of antibiotic resistance amongst bacterial pathogens, plant polyphenols are promising therapeutic options to combat bacterial infections.

Sorghum (*Sorghum bicolor* (L.) Monench) is a dryland cereal crop that produces bioactive polyphenols and performs well in arid climates, which is critical for sustainability in a continuously warming climate. Sorghum polyphenols have demonstrated antioxidant and anticancer activities [3,4]. Moreover, sorghum polyphenols have the ability to alleviate symptoms of colitis in addition to restoring microbiome diversity [9,10]. Despite exploration into the health benefits and bioactivity of sorghum polyphenols, their antimicrobial potential has not been explored.

*Legionella pneumophila* is facultative intracellular bacterial pathogen that naturally inhabits freshwater environments where it parasitizes and replicates within unicellular protozoa [11]. Inhalation of *Legionella*-contaminated aerosols from anthropomorphic fresh-water environments causes severe inflammatory pneumonia in immunocompromised individuals called Legionnaires’ Disease through uncontrolled bacterial replication within alveolar macrophages. To replicate within eukaryotic phagocytes, *L. pneumophila* employs a Dot/Icm type IV secretion system, which translocates hundreds of bacterial effector protein virulence factors directly into infected host cells [12]. Effector translocation by the Dot/Icm secretion system is essential for biogenesis of *L. pneumophila*’s replicative niche, the *Legionella* containing vacuole (LCV) and intracellular replication.

As an accidental pathogen of humans, *L. pneumophila* does not transmit between individuals and is highly susceptible to mammalian innate immune defenses. Specifically, *L. pneumophila* replication in macrophage is potently attenuated by proinflammatory cytokines, such as interferon (IFN)-γ and tumor necrosis factor (TNF) [13–17]. A previous study revealed that *L. pneumophila* is restricted in mouse macrophages by EGCg via increased inflammatory cytokine production [7]. The impact of EGCg on the macrophages was responsible for bacterial restriction since *L. pneumophila in vitro* replication was unaffected [7]. In the present study, we evaluated the impact of high-polyphenol extracts (HPE) from sorghum on *L. pneumophila* replication within macrophages. We discovered that HPE-treated macrophages were restrictive to *L. pneumophila*, which was not a consequence of macrophage cell death or direct antimicrobial restriction of *L. pneumophila in vitro*. Moreover, unlike tea catechin-mediated restriction, TNF secretion from *L. pneumophila*-infected macrophages was decreased following HPE treatment, which suggests that sorghum polyphenols elicit their antimicrobial activity through a novel mechanism.

## Materials and Methods

### Bacterial strains, tissue culture, reagents and growth conditions

*Legionella pneumophila* Philadelphia-1 SRS43 wild-type, Δ*flaA* and *dotA*::Tn [18,19] were cultured on supplemented charcoal–N (2-acetamido)-2-aminoethanesulfonic acid (ACES)-buffered yeast extract (CYE) and grown at 37°C as described previously [18]. Isolated colonies from CYE agar plates were used to generate 48 h heavy patches of bacteria, which were then used for infection. Liquid cultures were grown at 37°C in supplemented ACES-buffered yeast extract (AYE) as described [20,21].

RAW 264.7 cells (a gift from Dr. Craig Roy, Yale University) were maintained at 37°C/5% CO_2_ in Dulbecco’s Modified Eagle Medium (DMEM; Gibco) supplemented with 10% heat-inactivated fetal bovine serum (HIFBS; Gibco). For infections and assays, cells were seeded and infected in low-serum media (2.5 % HIFBS) where indicated. RAW cells were used from passages 4 to 15. All chemicals were purchased from MilliporeSigma (St. Louis, MO) unless otherwise specified.

### Generation of high-polyphenol sorghum bran extracts (HPE)

Sorghum accession PI570481 (tall and photo-period sensitive germplasm of Sudan origin) was obtained from Kansas State University, Agricultural Research Center, Hays, Kansas for its high phenolic content and previously demonstrated anti-cancer activity in tissue culture models and used for evaluation in this study [3]. Total sorghum phenolic extract was performed as previously described [22]. In brief, sorghum was decorticated, and bran was ground in-house. Bran powder (10% w/v) was suspended in extraction solvent, consisting of 70% v/v ethanol with 5% w/v citric acid. The samples were placed on a shaker at room temperature (20°C) for 2 h and then stored at −20°C overnight. The following day, the sample was centrifuged at 3000 r.c.f. for 10 min. The supernatant was collected as total phenolic extract and the pellet was discarded.

### Quantification of *L. pneumophila* replication *in vitro*

*Legionella pneumophila* was cultured on CYE agar and 48 h heavy patches were used to subculture bacteria into supplemented AYE media (see above). All cultures were diluted to an OD_600_ of 0.5 and were grown in triplicates at 37°C with shaking. HPE-treated cultures were grown in the presence of 1.25 mg/mL HPE and vehicle control cultures were grown in a volume equivalent of vehicle [5% citric acid (w/v), 70% ethanol (v/v)]. At the indicated time points, OD_600_ was measured and bacteria were plated on CYE agar for colony forming unit (CFU) enumeration.

### Infection of RAW 264.7 cells with *Legionella pneumophila*

RAW 264.7 cells were seeded in 24-well plates at 2 x 10^5^ cells/well in low-serum media (DMEM + 2.5% HI FBS) one day prior to infection. Cells were infected with *L. pneumophila* at a multiplicity of infection (MOI) of 1 as described previously [19] in the presence of HPE or vehicle as indicated. Plates were centrifuged at 250 r.c.f. to synchronize infection and incubated at 37°C/5% CO2 for 1 h. Media were aspirated and cells were washed gently 3 times with PBS and media were replaced with the same concentrations of HPE or vehicle. At the indicated time points, cells were lysed in sterile ultrapure water and bacteria were plated on CYE agar for CFU enumeration. Data are presented at either CFU per well at the indicated time point or fold replication at 48 h post-infection (normalization of CFU counts to counts at 1 h post-infection; day 0).

### Cytotoxicity assay

To determine if sorghum HPE treatment impacted viability of RAW cells, cytotoxicity was quantified by lactate dehydrogenase (LDH) release assay using the Promega CytoTox™ 96 Non-Radioactive Cytotoxicity Assay according to manufacturers’ instructions. Briefly, RAW cells were seeded at 2.5 x 10^5^/well in 24-well tissue culture plates in low-serum media one day prior to infection. Cells were infected at a MOI of 1 in the presence of 0.652 mg/mL HPE, 1.25 mg/mL HPE or volume equivalent of vehicle (1.25 μL/mL) for 1 h. Media were aspirated and cells were washed 3 times with sterile PBS followed by replenishing media in the presence or absence of HPE or vehicle. Lysis solution (1X) was added to control wells one hour prior to collecting supernatants. At 6 h post-infection, plates were centrifuged at 250 r.c.f. and supernatants were transferred to a sterile 96-well plate followed by addition of CytoTox™ reagent and incubation at room temperature in the dark for 30 min. Stop solution was added and absorbance was measured at 490 nm in a Victor 2 microplate reader (PerkinElmer). Background absorbance values were subtracted and % cytotoxicity was calculated by normalizing data to lysis buffer control (100% cytotoxicity).

### Caspase 3 cleavage assay

RAW 264.7 cells were seeded at 1 x 10^6^ in low-serum media in 6-well tissue culture plates one day prior to infection. Cells were treated with 1.25 mg/mL HPE or volume equivalent of vehicle and either infected with *L. pneumophila* at a MOI of 1 or left uninfected. As a control for induction of apoptosis, cells were treated with 10 μM staurosporine for 3 h [23]. Cells were treated and/or infected for a total of 4 h followed by lysis in 120 μL of ice-cold RIPA buffer [10 mM Tris,(pH 7.5), 100 mM NaCl, 1 mM EDTA, 1% NP40 (v/v), 0.1% SDS (v/v), 0.5% sodium deoxycholate (w/v)] supplemented with fresh 1X ProBlock Gold Mammalian Protease Inhibitor Cocktail and 1X EDTA (GoldBio). Cells were scraped into pre-chilled 1.5 mL microcentrifuge tubes and lysates were clarified by centrifugation at 14,000 r.c.f. at 4°C for 10 min. Clarified lysates (60 μL) were added to 3X Laemmli sample buffer (30 μL) and boiled for 10 min followed by SDS-PAGE and Western blot analysis.

### SDS-PAGE and Western Blot

Boiled protein samples were loaded onto 12% SDS-PAGE gels and separated by electrophoresis. Proteins were transferred to polyvinylidene fluoride (PVDF) membranes using a BioRad wet transfer cell. Membranes were blocked in blocking buffer [5% non-fat milk powder dissolved in tris-buffered saline, 0.1% Tween-20 (TBST)]. Primary antibodies [α-caspase 3 (#9662S; Cell Signaling Technology), α-β-actin (#4967S; Cell Signaling Technology)] were diluted in blocking buffer (1:1000) and incubated with membranes at 4°C overnight with rocking. Membranes were washed in TBST 3X 10 min with rocking at room-temperature. Horseradish peroxidase (HRP)-conjugated secondary antibodies (goat α-mouse-HRP or goat α-rabbit-HRP; ThermoFisher) were diluted to 1:5000 in blocking buffer and incubated with membranes for 1-2 h at room temperature with rocking. Membranes were washed in TBST as above. Membranes were incubated with ECL substrate (GE Amersham) and imaged by chemiluminescence using an Azure Biosystems c300 Darkroom Replacer. For loading control (β-actin) blots, membranes were stripped with Thermo Scientific Restore Western Blot Stripping Buffer (ThermoFisher) according to manufacturers’ instructions.

### Enzyme-linked immunosorbent assay (ELISA)

RAW 264.7 cells were seeded in 24-well tissue culture plates at 2.5 x 10^5^ in low-serum media one day prior to infection. Cells were infected with *L. pneumophila* at a MOI of 30 and either treated with the indicated concentrations of HPE or volume equivalent of vehicle. One hour post-infected, media were aspirated and cells were washed 3X with sterile PBS. Media and HPE/vehicle treatments were replaced and cells were incubated for an additional 3 h. Lysates were harvested from cells and stored at −20 °C until use. TNF was quantified using the Mouse TNF-α ELISA MAX™ Deluxe Kit (BioLegend) according to manufacturers’ instructions.

### Statistical analysis

Statistical analysis was performed with GraphPad Prism software using unpaired Students *t*-test with a 95% confidence interval. For all experiments, data are presented as mean ± standard deviation of samples in triplicates.

## Results

### High-polyphenol extracts (HPE) from *Sorghum bicolor* restrict *Legionella pneumophila* intracellular replication within RAW 264.7 cells

The polyphenol epigallocatchenin gallate (EGCg) from tea (*Camellia sinensis*) enhances restriction of *L. pneumophila* intracellular replication within mouse macrophages [6,7]. Thus, we evaluated whether *S. bicolor* high-polyphenol extracts (HPE) would also be sufficient to restrict *L. pneumophila* intracellular replication. Flagellin (FlaA) from *L. pneumophila* activates the NLRC4 inflammasome, which results in bacterial restriction in mouse macrophages [24,25]. To circumvent this issue, we utilized a flagellin deficient (Δ*flaA*) *L. pneumophila* strain for infection experiments. Replication of *L. pneumophila* Δ*flaA* within mouse RAW 264.7 macrophages was quantified in the presence or absence of HPE. Initially, cells were treated with 1.25 mg/mL HPE or volume equivalent of vehicle (see *Materials and Methods*) and *L. pneumophila* intracellular replication was quantified by enumeration of colony forming units (CFU) for up to 48 h post-infection. Replication of *L. pneumophila* was significantly decreased within RAW cells treated with HPE compared to vehicle control (**Fig 1A**, \**P*<0.05, \*\**P*<0.01). To determine if growth attenuation by HPE was dose-dependent, we treated RAW 264.7 cells with increasing concentrations of HPE (0.625 mg/mL and 1.25 mg/mL) or vehicle control and quantified *L. pneumophila* intracellular replication. Compared to vehicle-treated cells, *L. pneumophila* intracellular replication with significantly attenuated in macrophages treated with 0.625 mg/mL (*P*<0.05, **Fig 1B**)and further decreased within cells treated with 1.25 mg/mL HPE (*P*<0.01, **Fig 1B**). Together, these data suggest that sorghum HPE treatment is sufficient to restrict *L. pneumophila* intracellular replication in a dose dependent manner.

**Figure 1.**
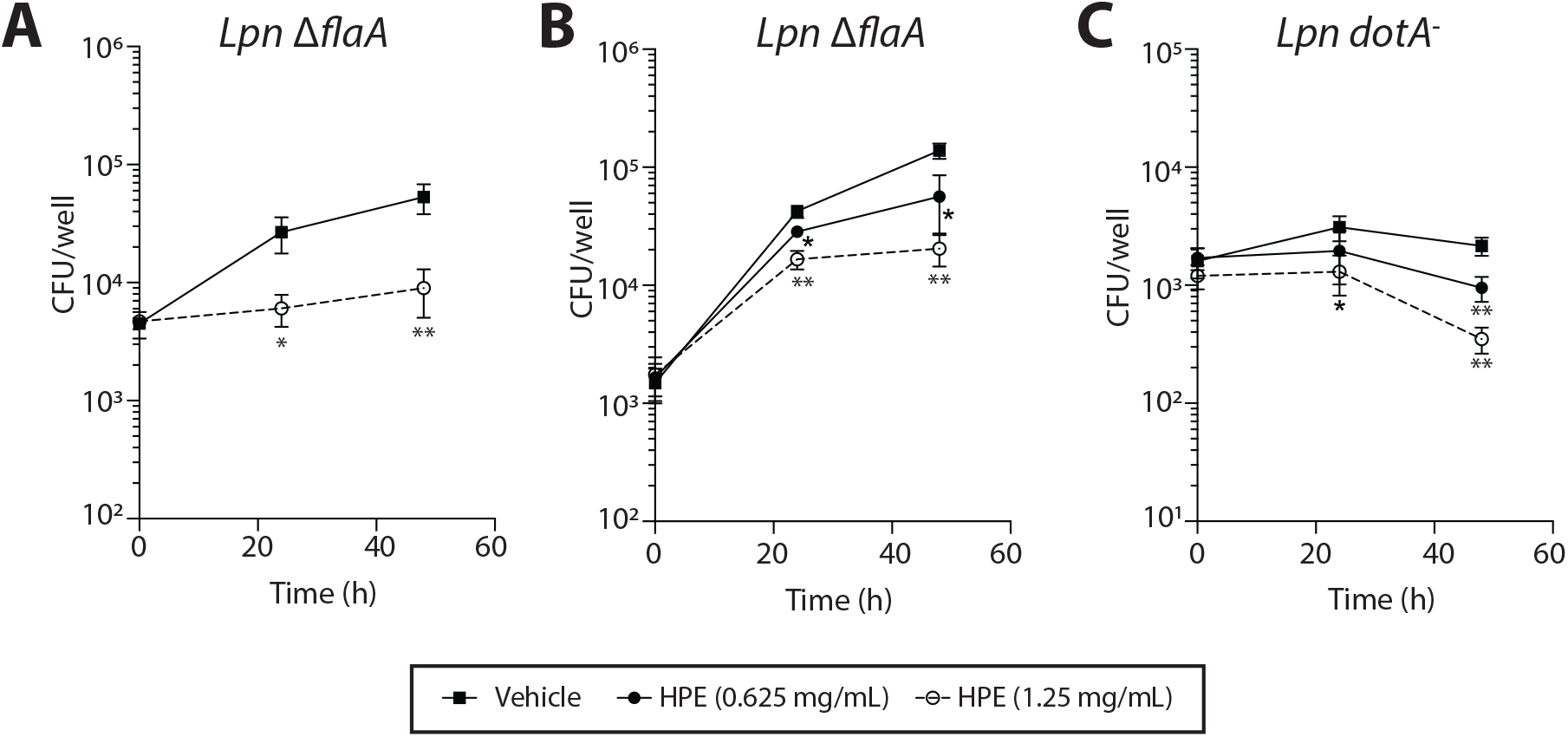
High-polyphenol sorghum extract (HPE) enhances bacterial killing by RAW 264.7 macrophages. **(A)** Quantification *L. pneumophila* Δ*flaA* recovered from RAW 264.7 macrophages treated with either HPE (1.25 mg/mL) or vehicle control. **(B)** Quantification *L. pneumophila* Δ*flaA* recovered from RAW 264.7 macrophages treated with either 1.25 mg/mL, 0.625 mg/mL or volume equivalent vehicle. **(C)** Quantification *L. pneumophila dotA*::Tn recovered from RAW 264.7 macrophages treated with either 1.25 mg/mL, 0.625 mg/mL or volume equivalent vehicle. Colony forming units (CFU) were enumerated at the indicated times post-infection. Data are presented as mean ± standard deviation (s.d.) of samples in triplicates. Asterisks denote statistical significance (\*\**P*<0.01, \**P*<0.05) by Students *t*-test compared to vehicle control. Data are representative of at least two independent experiments.

We subsequently evaluated whether HPE-mediated growth defects were restricted to replicating bacteria. RAW264.7 cells were treated with either vehicle control, 0.625 mg/mL, or 1.25 mg/mL HPE and infected with a strain of *L. pneumophila* with a loss-of-function mutation in *dotA* (*dotA*::Tn), which encodes a component of the Dot/Icm secretion system that is essential for effector translocation and thus bacterial intracellular replication. Although not replicating, the *dotA*::Tn strain persists within RAW 264.7 treated with vehicle control up to 48 h; however, the this strain was cleared more rapidly from RAW 264.7 cells treated with HPE in a dose-dependent manner (**Fig 1C**). These data suggest that HPE-treatment enhances the microbicidal activity of RAW 264.7 cells.

### *Legionella pneumophila* replication *in vitro* is not attenuated by sorghum HPE

We subsequently evaluated whether HPE has antimicrobial activity against *L. pneumophila in vitro*. Plate-grown *L. pneumophila* were sub-cultured into rich liquid medium in the presence of either 1.25 mg/mL HPE or vehicle control and *L. pneumophila* was quantified over 24 h by OD_600_ measurement or quantification of CFU (see *Materials and Methods*). Addition of 1.25 mg/mL HPE to rich medium did not impair *L. pneumophila* replication compared to vehicle control (**Fig 2**), suggesting that HPE is not microbicidal on its own. Thus, HPE-mediated restriction of *L. pneumophila* intracellular replication is likely not due to microbicidal activity of HPE.

**Figure 2.**
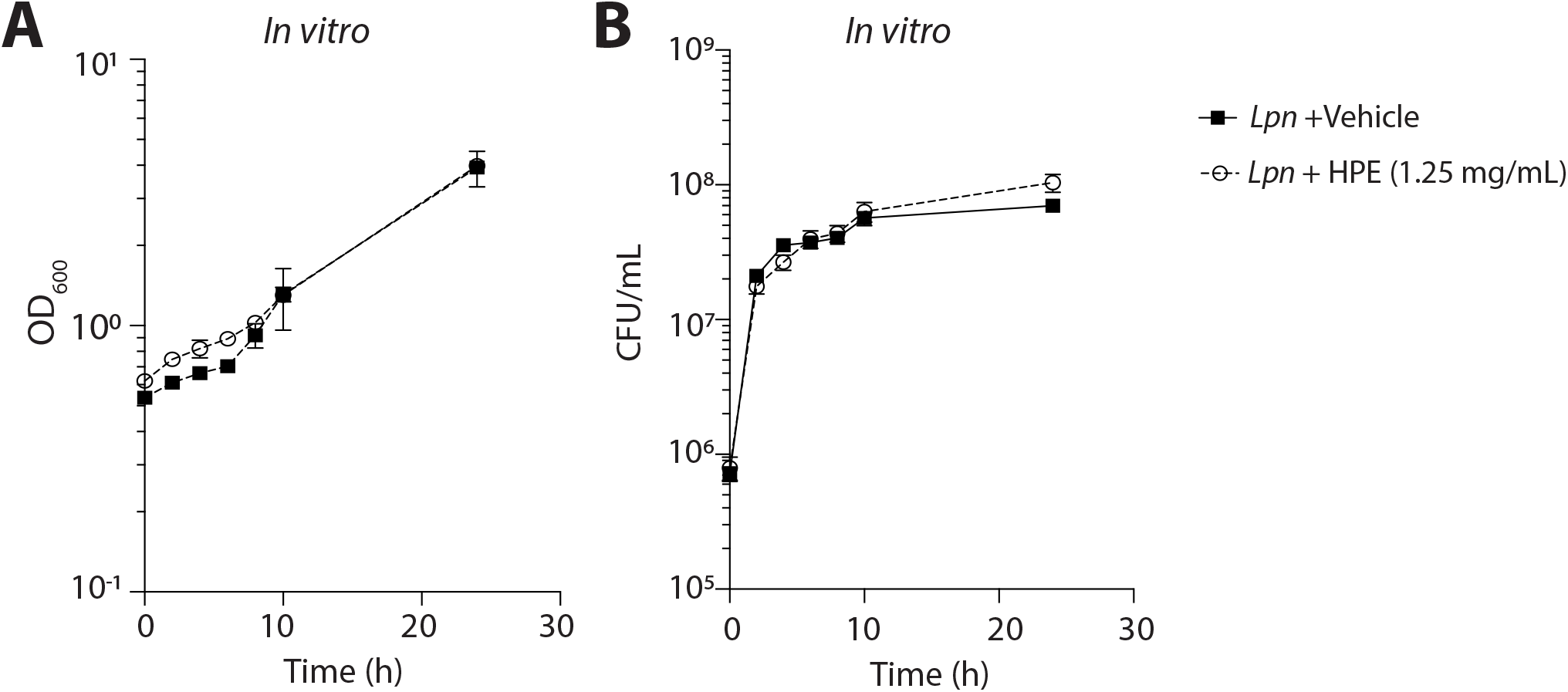
Sorghum HPE does not attenuate *L. pneumophila* replication *in vitro*. *L. pneumophila* was grown in the presence of 1.25 mg/mL HPE or volume equivalent of vehicle over 24 and **(A)** OD_600_ was measured or **(B)** CFU were quantified at the indicated times. Data are presented as mean ± s.d. of samples in triplicates. Data are representative of two independent experiments.

### Attenuation of *L. pneumophila* intracellular replication is not due to HPE-mediated macrophage cell death

HPE was shown to induce death in immortalized carcinoma cell lines. Thus, we evaluated whether restriction of *L. pneumophila* by HPE was due to death of RAW 264.7 cells. To determine whether HPE was toxic to RAW 264.7 cells, we utilized a lactate dehydrogenase (LDH) release assay, which is used to quantify cell lysis. To determine if HPE was toxic to RAW 264.7 cells, cells were treated with 1.25 mg/mL HPE or vehicle control and cytotoxicity was quantified by LDH release assay (see *Materials and Methods*). We observed low levels of cytotoxicity (< 2%) in both vehicle and HPE-treated cells (**Fig 3A**). To determine if RAW 264.7 cell viability in the presence of HPE was affected by *L. pneumophila* infection, cytotoxicity of HPE or vehicle treated cells infected with *L. pneumophila* was quantified. We found that there were no differences in viability of *L. pneumophila*-infected RAW 264.7 cells treated with HPE and vehicle treated cells (**Fig 3A**). Thus, HPE-mediated restriction of *L. pneumophila* intracellular replication is independent of host cell lysis.

**Figure 3.**
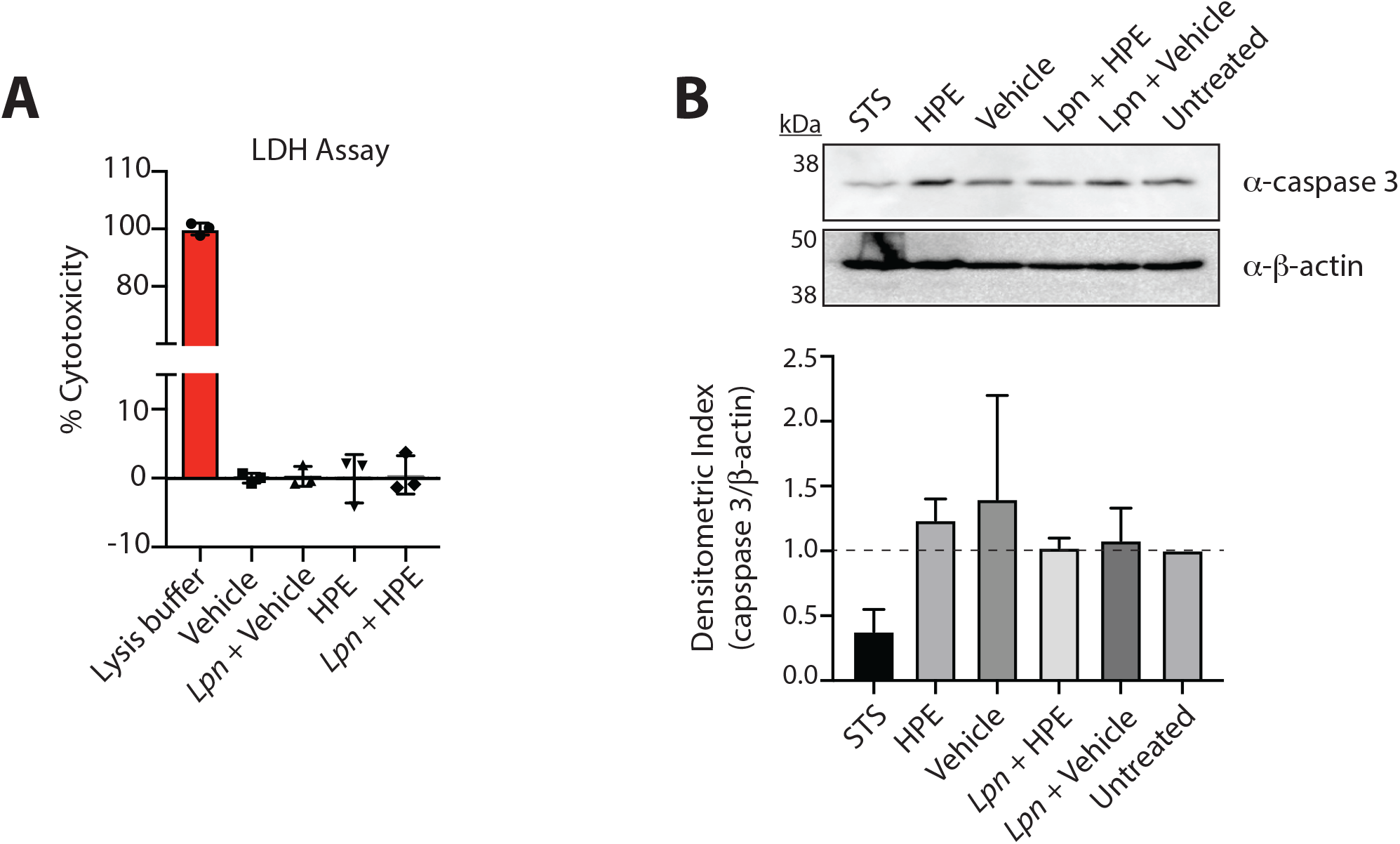
HPE does not impair RAW 264.7 cell viability. **(A)** RAW 264.7 cell viability was measured by LDH release assay after 4 h of treatment with 1.25 mg/mL HPE or volume equivalent of vehicle in the presence or absence of *L. pneumophila* Δ*flaA* infection (MOI of 1), as indicated. Lysis solution was used as a control for 100% cytotoxicity. **(B)** RAW 264.7 cells were treated with 1.25 mg/mL HPE or volume equivalent of vehicle in the presence or absence of *L. pneumophila* Δ*flaA* infection (MOI of 1), as indicated, and abundance of full-length caspase 3 was visualized by Western blotting (top panel) and quantified by densitometric analysis relative to the β-actin loading control (bottom panel). As a control for apoptosis, cells were treated with staurosporine (STS) for 3 h. Data shown are representative of two independent experiments.

Previous studies revealed that HPE can cause apoptosis of tumor cells. Initial stages of apoptosis are not accompanied by cell lysis and thus would not be detectable by LDH release assay. Therefore, to determine if HPE treatment caused RAW 264.7 cells to undergo apoptosis, we used Western blot and densitometry to quantify full length caspase 3 compared to actin. We were unable to visualize the cleaved fragment of caspase 3; however, RAW 264.7 cells treated with staurosporine (STS), which induces apoptosis, contained decreased levels of full-length caspase 3 compared to untreated cells (**Fig 3B, C**), indicating that decreased abundance of full-length caspase 3 is evidence of apoptosis. Caspase 3 levels were unchanged in cells treated with HPE compared to the vehicle control and this was also not affected by *L. pneumophila* infection (**Fig 3C**). Thus, HPE-mediated apoptosis of RAW 264.7 cells is likely not the reason for decreased *L. pneumophila* intracellular replication.

### HPE treatment impairs tumor necrosis factor (TNF) secretion from *L. pneumophila*-infected macrophages

Previous work demonstrated that EGCg increased TNF secretion from *L. pneumophila*-infected macrophages [7]. We therefore hypothesized that sorghum polyphenols may also augment TNF secretion by *L. pneumophila*-infected macrophages. To test this hypothesis, we quantified TNF secretion from macrophages infected with *L. pneumophila* for 4 h and treated with the indicated concentrations of HPE or vehicle control. Interestingly, we found that treatment with HPE resulted in a significant decrease in TNF production from infected macrophages (**Fig 4**). Thus, unlike EGCg, sorghum polyphenols decreased TNF production by macrophages infected with *L. pneumophila*.

**Figure 4.**
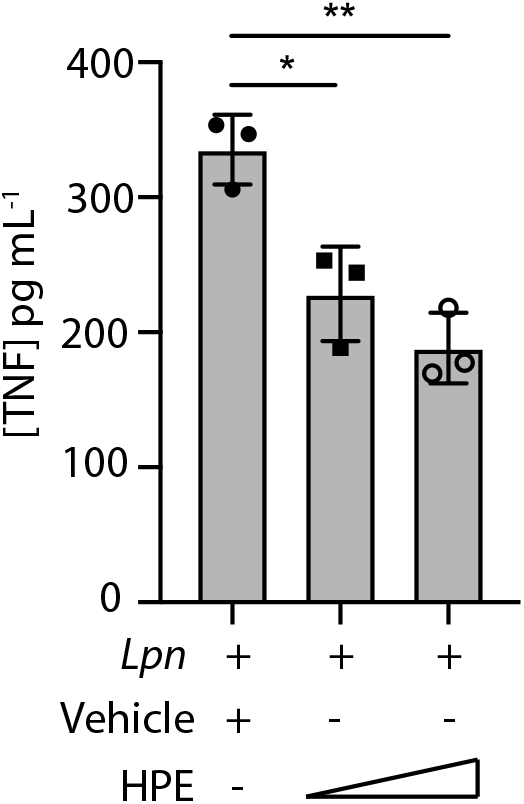
HPE impairs TNF production by *L. pneumophila*-infected macrophages. ELISA to quantify TNF produced by RAW cells infected with *L. pneumophila* Δ*flaA* (MOI of 30) and treated with 0.625 mg/mL, 1.25 mg/mL HPE or vehicle for 4 h. Asterisks indicate statistical significance by Students’ *t*-test (\**P*<0.05, \*\**P*<0.01). Data are shown as mean ± s.d. of samples in triplicates and are representative of at least two independent experiments.

## Discussion

In this study, we revealed that polyphenols from sorghum bran are capable of increasing macrophage restriction of *L. pneumophila* in culture. High-polyphenol extracts (HPE) attenuated *L. pneumophila* replication in a dose-dependent manner within macrophages but not in rich media *in vitro*. HPE treatment also resulted in enhanced clearance of the avirulent *dotA*::Tn strain, suggesting that HPE-treated macrophages actively eliminate intracellular *L. pneumophila* as opposed to stalling replication. Moreover, we found that HPE attenuates TNF secretion during *L. pneumophila* infection, suggesting that sorghum polyphenols function by a mechanism distinct from EGCg. Together, these data suggest that HPE promotes bacterial killing by macrophages through a novel mechanism.

Sorghum polyphenols comprise a family of compounds with diverse chemical structures and biological activities. The major components of sorghum polyphenols are tannins, phenolic acids and flavonoids, all of which have documented health benefits [26]. Our work demonstrates a role for polyphenols in bacterial restriction, but the specific component(s) of the extract responsible for this activity is unknown. Since EGCg is a flavonoid, it is possible that sorghum contains analogous flavonoids that function similarly since several sorghum genotypes produce relatively high levels of flavonoids compared to other crops [27]. However, the component(s) of sorghum polyphenol extracts responsible for enhanced macrophage killing will be the subject of future investigation.

Sorghum polyphenols potently inhibit growth of cancer cells through generation of reactive oxygen species (ROS), apoptosis and cell cycle arrest [3]. Viability of HT-29, HepG2 and Caco2 cells was significantly decreased following treatment with sorghum polyphenols [3,28]. Sorghum polyphenols were also found to dampen symptoms of colitis in a mouse model of disease, indicating potential to alleviate colitis and potentially prevent colitis-associated cancer [10]. Interestingly, sorghum HPE did not cause apoptosis or necrosis of RAW cells, which were derived from a murine leukemia virus-induced tumor [29]. Thus, sorghum polyphenols have distinct effects on epithelial cells and macrophages.

The antimicrobial properties of non-sorghum polyphenols have been documented *in vitro* and *in vivo*. The majority of these studies have focused on food-borne pathogens, but reveal the potential for natural and sustainable sources of antimicrobial compounds. While in vitro restriction of many bacterial, fungal and viral pathogens has been evidenced (reviewed in [1,30,31]), polyphenols are also capable of specific inhibition of pathogen virulence. A striking example of this is the ability of tannic acid and *n*-propyl gallate to protect mice from *Helicobacter pylori*-mediated disease [32]. Ruggiero *et al*. discovered that administration of polyphenols to mice in drinking water decreased *H. pylori*-mediated gastritis and bacterial load in the stomachs of mice. Polyphenols from red wine and tea were also sufficient to inhibit the function of the *H. pylori* vacuolating toxin (VacA), demonstrating specific anti-virulence properties for these compounds [32,33]. Moreover, infectivity and replication of the obligate intracellular pathogen *Chlamydia pneumoniae* was also inhibited by diverse plant polyphenols [34].

Plant polyphenols have been demonstrated to modulate mammalian immunity (reviewed in [35]). Namely, polyphenols have anti-inflammatory and antioxidant activity and our work demonstrates that sorghum HPE similarly limits production of the pro-inflammatory cytokine, TNF during *L. pneumophila* infection. These data suggest that HPE-mediated microbicidal activity of macrophages is independent of TNF. Non-inflammatory macrophage restriction of *L. pneumophila* by HPE is very intriguing and suggests potential for use of sorghum polyphenols as antimicrobial factors in immunosuppressed individuals that are incapable of mounting a robust inflammatory response to infection. Based on our observation that sorghum HPE increases microbicidal activity of macrophages, sorghum HPE may stimulate increased lysosomal fusion to the LCV.

Ultimately, our work provides the foundation for investigation into how sorghum polyphenols influence macrophage physiology to combat bacterial infection. The mechanism by which sorghum HPE attenuates intracellular replication and whether sorghum HPE restrict *L. pneumophila* in a mouse model of infection will be the subject of future studies.

## Acknowledgements

This work was funded by U.S. Department of Agriculture Research Service project # 3020-43440-001-00D and institutional start-up funds from Kansas State University (to SRS).

